# Ethograms reveal a fear conditioned visual cue to organize diverse behaviors

**DOI:** 10.1101/2024.09.10.612214

**Authors:** David C. Williams, Amanda Chu, Nicholas T. Gordon, Aleah M. DuBois, Suhui Qian, Genevieve Valvo, Selena Shen, Jacob B. Boyce, Anaise C. Fitzpatrick, Mahsa Moaddab, Emma L. Russell, Liliuokalani Counsman, Michael A. McDannald

## Abstract

Recognizing and responding to threat cues is essential to survival. In rats, freezing is the most common behavior measured. Previously we demonstrated a threat cue can organize diverse behaviors (Chu et al., 2024). However, the experimental design of Chu et al. (2024) was complex and the findings descriptive. Here, we gave female and male Long Evans rats simple paired or unpaired presentations of a light and foot shock (8 total) in a conditioned suppression setting, using a range of shock intensities (0.15, 0.25, 0.35 or 0.5 mA). We found that conditioned suppression was only observed at higher foot shock intensities (0.35 mA and 0.5 mA). We constructed comprehensive, temporal ethograms by scoring 22,272 frames of behavior for 12 mutually exclusive behavior categories in 200 ms intervals around cue presentation. A 0.5 mA and 0.35 mA shock-paired visual cue suppressed reward seeking, rearing and scaling, as well as light-directed rearing and light-directed scaling. The shock-paired visual paired cue further elicited locomotion and freezing. Linear discriminant analyses showed that ethogram data could accurately classify rats into paired and unpaired groups. Considering the complete ethogram data produced superior classification than considering subsets of behaviors. The results demonstrate diverse threat-elicited behaviors – in a simple Pavlovian fear conditioning design – containing sufficient information to distinguish the fear learning status of individual rats.

## Introduction

In a typical Pavlovian fear conditioning procedure, a neutral cue is paired with foot shock. As a result, the shock-paired cue is endowed with a multitude of properties (McDannald, 2023). One property is the ability to elicit behavior. Most well-known is that a shock-paired cue (or context) can elicit freezing (Bolles and Collier, 1976). Pavlovian conditioning procedures measuring freezing have been used to great effect to reveal neural bases of fear (LeDoux et al., 1988; De Oca et al., 1998; Goosens and Maren, 2001).

A shock-paired cue can suppress reward seeking (Estes and Skinner, 1941), commonly referred to as conditioned suppression. A simple explanation for conditioned suppression is response competition via freezing. A shock-paired cue that elicits freezing will suppress responding for reward. Indeed, conditioned freezing and conditioned suppress can be correlated in some (Bouton and Bolles, 1980; Mast et al., 1982) but not all (Chu et al., 2024) settings. However, neural manipulations that abolish freezing can leave conditioned suppression intact (Amorapanth et al., 1999; Lee et al., 2005; McDannald, 2010; McDannald and Galarce, 2011). Observing conditioned suppression in the absence of conditioned freezing means either non-freezing behaviors compete to inhibit reward seeking, or suppression is not achieved through direct behavioral competition.

We recently demonstrated that a shock-paired auditory cue will suppress reward seeking, elicit locomotion, rearing and jumping, and also elicit freezing (Chu et al., 2024). Though consistent with some studies, observing such diversity of threat-elicited behaviors is not common. It is possible we observed this diversity was due to the complexity of our experimental design. Rats discriminated between two or three cues associated with unique foot shock probabilities (i.e., danger, uncertainty, and safety). Rats also received many conditioning sessions. Further, the results of the prior study were descriptive. Percent behavior to danger and safety were measured for each category of interest then statistically compared. Behaviors specifically engaged by the danger cue were then determined. If there is a robust relationship between a cue’s learning status (pairing with shock) and these diverse behaviors, then we should be able to predict an individual’s learning status based on individual’s total behavior.

The primary goal of the present experiments was to comprehensively quantify behaviors controlled by a shock-paired visual cue. We chose a visual cue for three reasons. First, to test how well our auditory findings (Chu et al., 2024) generalize to another modality. Second, it is known that a visual cue supports conditioned suppression that is equivalent to an auditory cue, despite producing little conditioned freezing (Kim et al., 1996). Yet, in the study by Kim and colleagues non-freezing behaviors evoked by the shock-paired visual cue could not be established. This means that there is a small but important gap in the literature concerning visually-evoked, fear conditioned behaviors. Third, conditioned suppression following lesion-induced abolishment of conditioned freezing was obtained with visual cues (McDannald, 2010; McDannald and Galarce, 2011). If conditioned suppression is achieved through alternative behaviors these behaviors should be most readily observed in conditioned suppression setting using a visual cue.

A secondary goal asked more specifically if a shock-paired visual cue supports locomotion (also termed darting or flight). This goal is generally motivated by observations that flight responses can be readily obtained in Pavlovian fear conditioning settings (Gruene et al., 2015; Totty et al., 2021; Mitchell et al., 2022; Le et al., 2023; Borkar et al., 2024) but see (Trott et al., 2022). Specific motivation comes from our prior study (Chu et al., 2024) in which we observed locomotion elicited by a shock-paired auditory cue during conditioning sessions in which foot shock was present and extinction in which foot shock was absent.

To reveal behaviors engaged by a shock-paired visual cue we performed two experiments. Mildly food-deprived, Long Evans rats were trained to nose poke for food, then received Pavlovian fear conditioning over a baseline of rewarded poking. Both experiments used a between-subjects, 2 x 2 design (group x intensity) in which female and male rats (50/50 split) had a 10-s visual cue paired or unpaired with foot shock. In the vein of Holland (1979), we wanted to determine the lowest shock intensity that would support complete conditioned suppression (Holland, 1979). In Experiment 1 paired or unpaired rats received 0.15 or 0.25 mA foot shock. In Experiment 2 paired or unpaired rats received 0.35 or 0.5 mA foot shock. All rats received 8 total cue illuminations and shock presentations; simper and shorter experiments compared to our prior work (Chu et al., 2024).

TTL-triggered cameras were installed in testing chambers and were programmed to capture frames at 200-ms intervals prior to, during, and following cue light illumination. For Experiment 2, 22,272 frames were hand scored for twelve behaviors under categories of Immobility (freezing and stretching), Horizontal movement (locomotion and backpedaling), Vertical movement (rearing, scaling, and jumping) and Reward-related behavior (nose port and food cup) behavior. We further differentiated light-directed variants of each Vertical movement type. Comprehensive ethograms were constructed from the scored frames. Multivariate and univariate ANOVA performed on the ethogram data revealed behaviors that differed between groups. Latent discriminant analyses determined if group membership (paired vs. unpaired) could be classified from ethogram data.

## Results

Our first objective was to determine the lowest foot shock intensity that would support conditioned suppression to a visual cue. Rats were shaped to nose poke in a central port for a food reward which was delivered via a cup below. The cue light was positioned above the central port. Cue light presentation resulted in bright, focal illumination of the light itself as well as diffuse illumination of the entire context, which was otherwise dark (Figure 1A). In Experiment 1, rats received cue light presentations paired or unpaired with a 0.25 or 0.15 mA foot shock. In Experiment 2, rats received cue light presentations paired or unpaired with a 0.5 or 0.35 mA foot shock (Figure 1B). Behavior chambers were equipped with TTL-triggered cameras capturing 5 frames/s starting 5 s prior to cue light presentation and ending 2.5 s following cue offset.

**Figure 1.**
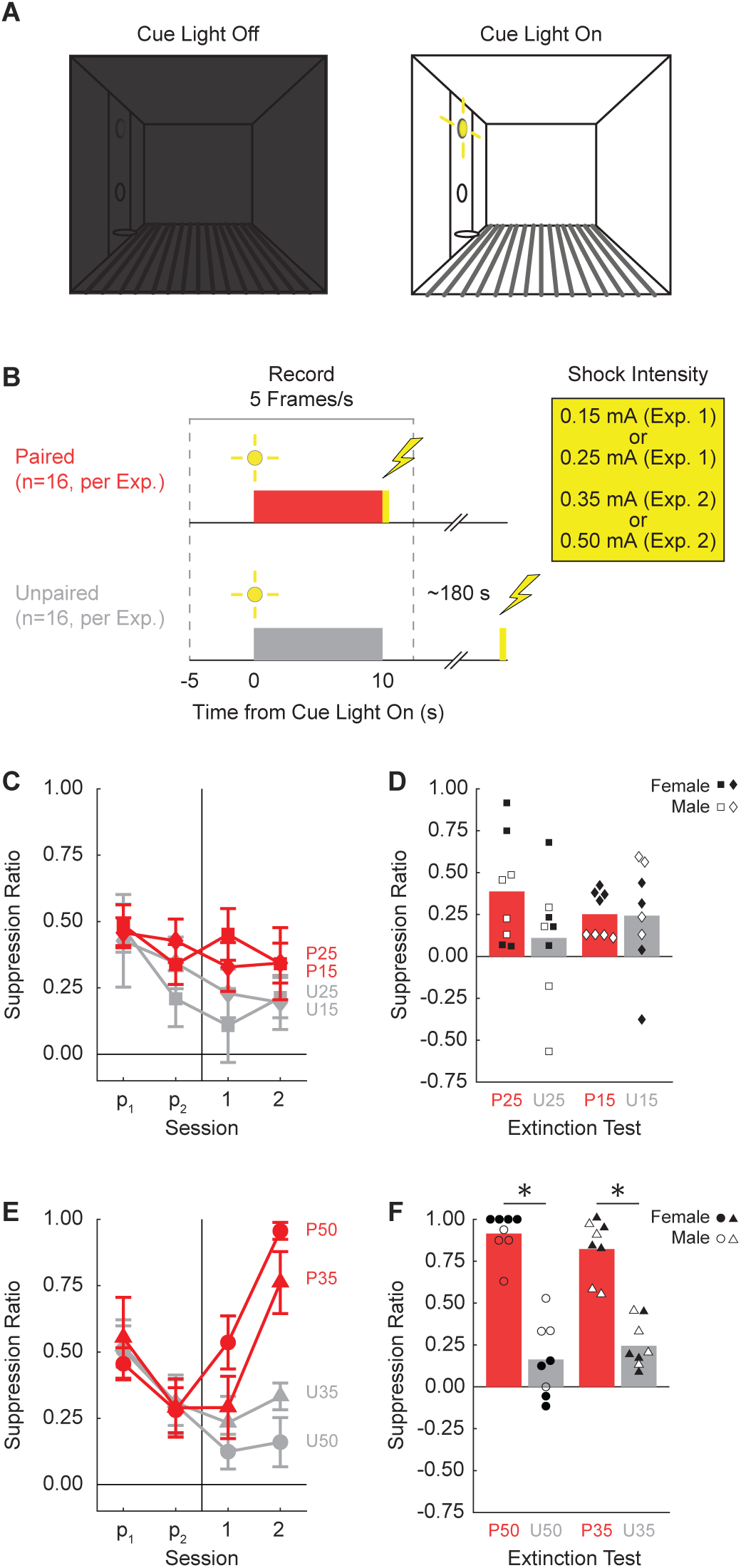
Experimental design and suppression ratio results. (**A**) Behavioral testing took place in a dark, Med Associates chamber equipped with a food cup, port, and cue light on one wall (left). Cue light illumination provided a discrete visual cue in addition to generally illuminating the chamber. (**B**) Rats in each experiment were divided into Paired (red; n=16) and Unpaired (gray; n=16) conditions. Experiment 1 used foot shock intensities of 0.15 mA and 0.25 mA, while Experiment 2 used foot shock intensities of 0.35 mA and 0.50 mA. (**C**) Mean ± SEM suppression ratio for Paired (red) and Unpaired (gray) rats receiving 0.25 mA (diamond) or 0.15 mA (square) foot shock over the two pre-exposure sessions (p1 and p2) and the two foot shock sessions (1 and 2) are shown for Experiment 1. Behavior frames were captured every 200 ms, 5 s prior to and 2.5 s following cue light presentation. (**D**) Group mean (bar) and individual mean (point) suppression ratio for the extinction test are shown for female (closed points) and male (open points), with color and shape meaning from C maintained. Experiment 2 suppression ratio data are shown in **E** and **F**. Formatting other than foot shock intensity (0.50 mA, circles; 0.35 mA, triangles) is identical to C and D.

### Higher, but not lower, shock intensities support conditioned suppression to a visual cue

Neither the 0.25 mA nor the 0.15 mA foot shock supported conditioned suppression to the visual cue. ANOVA for suppression ratio [within factors: session (4, 2 pre-exposure & 2 shock presentation); between factors: sex (female vs. male), group (paired vs. unpaired), and intensity (0.25 mA vs. 0.15 mA); Figure 1C] found no significant main effect of group (F_1,24_ = 3.15, *p* = 0.088), nor any significant group interactions (all Fs < 1, all *p*s > 0.5). In further support, ANOVA for the extinction session [within factors: trial (4); between factors: sex (female vs. male), group (paired vs. unpaired), and intensity (0.25 mA vs. 0.15 mA); Figure 1D] also found no significant main effect of group (F_1,24_ = 2.03, *p* = 0.17), nor any significant group interactions (all Fs < 2, all *p*s > 0.15). Two-tailed, independent samples t-tests found no significant differences between the paired and unpaired rats receiving the 0.25 mA foot shock (t = 1.60, *p* = 0.13), or the 0.15 mA foot shock (t = 0.06, *p* = 0.95).

By contrast, both the 0.5 mA and the 0.35 mA foot shock supported conditioned suppression to the visual cue. ANOVA for suppression ratio [within factors: session (4, 2 pre-exposure & 2 shock presentation); between factors: sex (female vs. male), group (paired vs. unpaired), and intensity (0.5 mA vs. 0.35 mA); Figure 1C] found a significant main effect of group (F_1,24_ = 13.5, *p* = 0.001), as well as a significant group x session interaction (F_3,72_ = 12.87, *p* = 8.03 x 10^-7^). No significant interactions with sex or intensity were observed (all Fs < 2.2, all *p*s > 0.1).

Conditioned suppression was apparent during extinction, with ANOVA [within factors: trial (4); between factors: sex (female vs. male), group (paired vs. unpaired), and intensity (0.5 mA vs. 0.35 mA); Figure 1D] revealing a significant main effect of group (F_1,24_ = 154.01, *p* = 6.21 x 10^-12^). Now, a significant sex x group interaction was observed (F_1,24_ = 9.21, *p* = 0.006). Paired rats showed higher suppression ratios than unpaired rats across sexes, but this difference was greater in Female rats. Two-tailed, independent samples t-tests found significant differences between the paired and unpaired rats receiving the 0.5 mA foot shock (t = 8.24, *p* = 3.52 x 10^-6^) and the 0.35 mA foot shock (t = 7.37, *p* = 9.68 x 10^-7^).

### Scoring 12 mutually exclusive behaviors with high inter-rater reliability and low inter-rater confusion

We defined (Table 1) and scored 12 mutually exclusive behaviors in 200-ms intervals around cue light presentation. These behaviors belonged to 4 different categories: Immobile (freezing and stretching), Reward (cup and port), Vertical (rearing, scaling, and jumping), and Horizontal (locomotion and backpedaling). Within the Vertical behavior we further separated rearing and scaling that were directed towards the light. Light-directed behavior reflect stimulus orienting while non-light-directed behavior reflects context exploration.

We focused our scoring efforts on Experiment 2, in which rats receiving the 0.5 mA and 0.35 mA foot shock intensities showed robust conditioned suppression. Six observers blind to group, sex, and intensity (see methods for blinding details) scored 22,272 total frames: 11,136 from the second foot-shock session, and 11,136 from the extinction session. Additionally, these six observers scored 696 frames spanning eight separate comparison trials. We determined inter-rater reliability by calculating the % identical observations for each observer-observer pair across the 696 frames (Figure 2A). Mean inter-rater reliability was 72.41%. Although inter-rater reliability was higher when fewer behaviors were present in a trial (e.g., 77.7% when 3 present); inter-rater reliability was 71.9% even when eight behaviors were present (Figure 2B).

**Figure 2.**
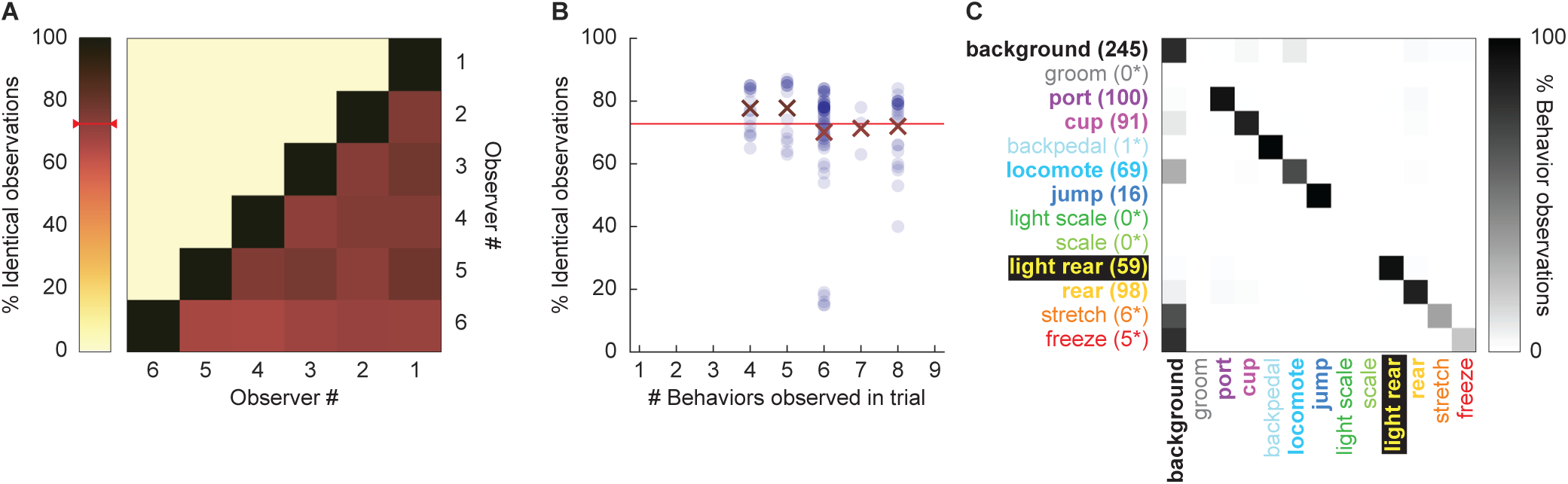
Hand scoring inter-rater reliability. (**A**) % Identical observations for the comparison frames was calculated for each observer-observer pair. Darker colors indicate higher % identical observations. The red bar on the scale shows mean % Identical observations across all observer-observer pairs. (**B**) % Identical observations is broken down by # of behaviors present during a trial, ranging from 4 to 8. Each individual point indicates a single observer-observer trial comparison. Large Xs indicate the mean % identical observations for all observer-observer comparisons for each # of behaviors. (**C**) A confusion matrix shows all observer-observer judgment pairs for the comparison trials. Y axis is observer 1 and x axis is observer 2. % Behavior observations is plotted by row with black indicating 100%, white 0% and shades of gray in between. The number of behavior observations is indicated for each in parentheses. Asterisks indicate behaviors for which fewer than 10 instances of the behavior were observed.

Disagreement rarely involved two observers assigning different, specific behaviors to a single frame. Instead, disagreement occurred when one observer assigned background to a frame, while the other observer assigned a specific behavior to that frame. To quantify disagreements, we constructed a confusion matrix (Figure 2C) composed of every single-frame judgment for every observer-observer pair. Zero confusion, identical observations for every frame, would manifest as a diagonal line across the matrix (top left to bottom right). Confusing a specific behavior and background would manifest as a horizontal line on the top edge or a vertical line on the left edge. Confusing a specific behavior for another specific behavior would manifest as values occurring between the diagonal and edges. The confusion matrix shows the vast majority of values fall on the diagonal, with most of the remaining values on the left edge. The confusion matrix results reveal high scoring specificity.

### Ethograms reveal diverse fear conditioned behaviors

We constructed comprehensive, high-temporal resolution ethograms for paired (Figure 3A) and unpaired (Figure 3B) rats from Experiment 2. For both groups, baseline behavior largely consisted of rearing, food cup and port behavior, with very small amounts of freezing and locomotion. Paired and unpaired behavior dramatically parted during cue light illumination. Paired rats increased freezing and locomotion during cue illumination, then increased locomotion, backpedaling and jumping during and following shock presentation. At the same time, paired rats decreased rearing and food cup behavior. By contrast, unpaired rats increased light scaling, rearing, and light rearing during cue light illumination. Although increases in light rearing were observed in paired rats, these increases were blunted compared to unpaired rats.

**Figure 3.**
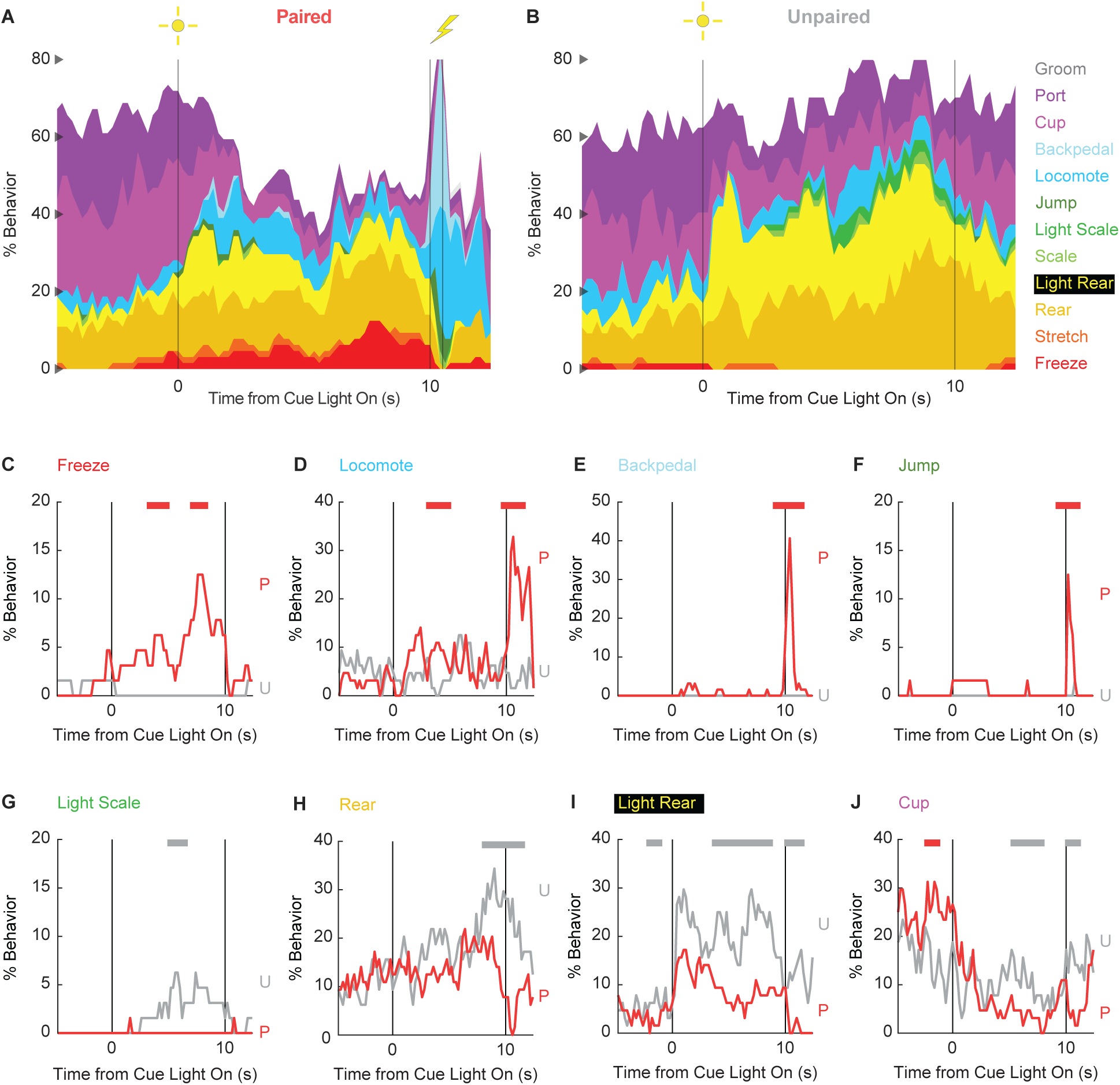
Conditioning ethograms. (**A**) Ethogram for paired rats shows % behavior (y axis) for each behavioral category in 200-ms intervals from 5-s prior to cue onset (time 0) to 2.5 s following cue offset (time 10). Paired rats had shock delivered at cue offset. Colors for each behavior category: Freeze (red), Stretch (orange), Rear (mustard), Light Rear (yellow), Scale (light green), Light Scale (green), Jump (dark green), Locomote (cyan), Backpedal (sky blue), Cup (magenta), Port (purple), and Groom (gray). (**B**) Ethogram for unpaired rats, details identical to paired rats. Unpaired rats did not receive shock at cue offset. Line graphs for the 8 behaviors showing a significant group x time interaction are shown in panels **C-J**. % Behavior is shown for paired rats (P, red) and unpaired rats (U, gray). X axis same as in A/B, y axis scaled to best visualize behavior patterns. Colored bars at the top of each axis indicate 1-s time periods in which paired and unpaired % behavior differed (independent samples t-test, *p* < 0.05). Red bars indicate greater behavior in paired rats while gray bars indicate greater behavior in unpaired rats.

In support of group differences across all behaviors, MANOVA for all 12 behaviors [factors: time (87, 200-ms bins), group (paired vs. unpaired), intensity (0.5 mA vs. 0.35 mA), and sex (female vs. male)] revealed a significant group x time interaction (F_1032,24768_ = 2.50, *p* = 7.41 x 10^-124^). MANOVA additional revealed significant group x time x sex (F_1032,24768_ = 1.13, *p* = 0.003) and group x time x intensity (F_1032,24768_ = 1.64, *p* = 5.47 x 10^-33^) interactions, which we return to in our univariate analyses.

To reveal behavior-specific differences between groups we performed univariate ANOVA using a Bonferroni-corrected *p*-value of 0.004167 (0.05/12) to reduce type 1 error. Univariate ANOVA [factors same as MANOVA] found significant group x time interactions for 8 of the 12 behaviors: freeze (F_86,2064_ = 2.13, *p* = *1.85 x 10^-8^*; Figure 3C), locomote (F_86,2064_ = 3.72, *p* = 5.04 x 10^-26^; Figure 3D), backpedal (F_86,2064_ = 14.95, *p* = 2.34 x 10^-159^; Figure 3E), jump (F_86,2064_ = 3.23, *p* = 2.98 x 10^-20^; Figure 3F), light scale (F_86,2064_ = 1.53, *p* = 0.003; Figure 3G), rear (F_86,2064_ = 1.75, *p* = 3.79 x 10^-5^; Figure 3H), light rear (F_86,2064_ = 1.47, *p* = 0.004; Figure 3I), and cup (F_86,2064_ = 2.05, *p* = 8.75 x 10^-8^; Figure 3J). The 8 behaviors differentially expressed by paired and unpaired rats varied in their direction and timing. Paired rats showed greater freezing only during the cue period, greater locomotion during the cue and post-cue periods, and greater backpedaling and jumping only during the post-cue period. Unpaired rats showed greater light scaling during only the cue period, but greater rearing, light rearing, and food cup behavior during the cue and post-cue periods.

Univariate ANOVA [factors same as above] further found significant group x time x intensity interactions for 3 of the 12 behaviors: locomote (F86,2064 = 1.52, *p* = 0.002), jump (F86,2064 = 2.58, *p* = 5.34 x 10^-13^), and backpedal (F86,2064 = 1.63, *p* = 2.64 x 10^-63^). Ethograms separating 0.5 and 0.35 mA rats are shown in Figure 3 – Supplemental Figure 1. Paired rats receiving the 0.5 mA foot shock jumped and locomoted following shock presentation, but rarely backpedaled. Unpaired rats receiving the 0.35 mA foot shock backpedaled then locomoted following shock presentation, but rarely jumped.

Finally, univariate ANOVA found significant group x time x sex interactions for 3 of the 12 behaviors: rear (F86,2064 = 1.80, *p* = 1.41 x 10^-5^), jump (F86,2064 = 1.93, *p* = 1.12 x 10^-6^), and backpedal (F86,2064 = 1.63, *p* = 2.79 x 10^-4^). Line graphs separately plotting female and male rats for each of the 3 behaviors are shown in Figure 3 – Supplemental Figure 2. Sex differences were not all or none – both paired females and males reduced rearing and increased jumping and backpedaling. Instead, differences amount to behavior timing and magnitude. Paired female rats suppressed rearing during late cue presentation while males suppressed post-cue. Paired female rats had a longer locomote bout, on average, compared to males following shock presentation. Paired males showed greater levels of backpedaling compared to females following shock presentation.

### Behavioral diversity persists in extinction

To determine if any behaviors were differentially expressed in the absence of foot shock, we constructed complete ethograms for paired (Figure 4A) and unpaired (Figure 4B) rats during extinction testing. As during conditioning, baseline behavior largely consisted of food-related behavior and rearing, and was similar for paired and unpaired rats. Paired and unpaired behavior parted following cue light illumination. Paired rats increased freezing and locomotion during cue illumination, then increased locomotion following cue offset. Freezing accounted for less than ∼5% of total behavior. Unpaired rats increased scaling throughout cue illumination and maintained high levels of light rearing throughout cue and post-cue periods. Both scaling and light rearing were suppressed in paired rats.

**Figure 4.**
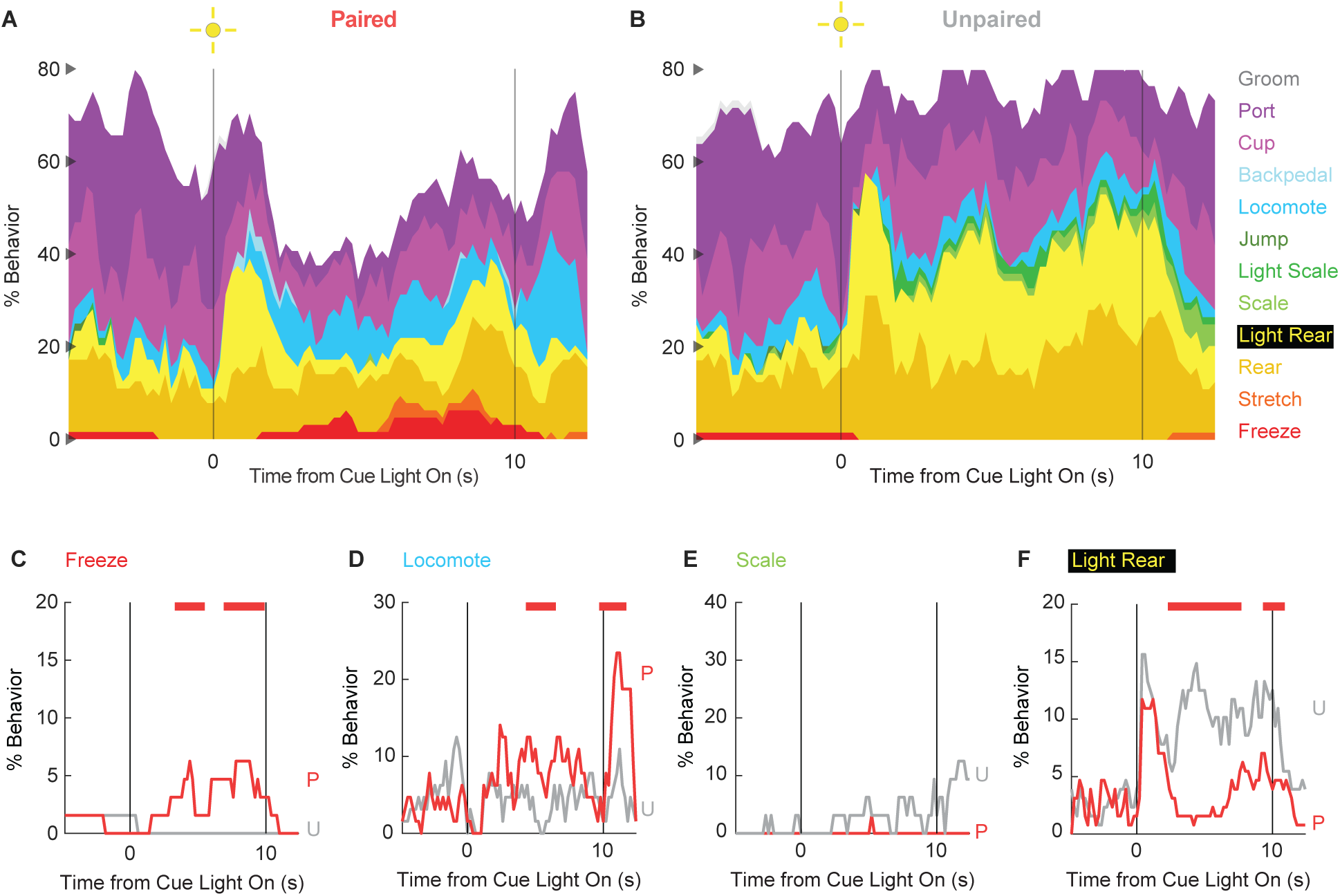
Extinction ethograms. (**A**) Ethogram for paired rats shows % behavior (y axis) for each behavioral category in 200-ms intervals from 5-s prior to cue onset (time 0) to 2.5 s following cue offset (time 10). No shock was delivered during this session. Colors for each behavior category: Freeze (red), Stretch (orange), Rear (mustard), Light Rear (yellow), Scale (light green), Light Scale (green), Jump (dark green), Locomote (cyan), Backpedal (sky blue), Cup (magenta), Port (purple), and Groom (gray). (**B**) Ethogram for unpaired rats, details identical to paired rats. Line graphs for the 4 behaviors showing a significant group x time interaction are shown in panels **C-F**. % Behavior is shown for paired rats (P, red) and unpaired rats (U, gray). X axis same as in A/B, y axis scaled to best visualize behavior patterns. Colored bars at the top of each axis indicate 1-s time periods in which paired and unpaired % behavior differed (independent samples t-test, *p* < 0.05). Red bars indicate greater % behavior in paired rats while gray bars indicate greater % behavior in unpaired rats.

In support of group differences across all behaviors, MANOVA for all 12 behaviors [factors: time (87, 200-ms bins), group (paired vs. unpaired), intensity (0.5 mA vs. 0.35 mA), and sex (female vs. male)] revealed a significant group x time interaction (F1032,24768 = 1.50, *p* = 4.80 x 10^-22^). MANOVA additional revealed significant group x time x intensity interaction (F1032,24768 = 1.21, *p* = 1.21 x 10^-6^) but no significant group x time x sex interaction (F1032,24768 = 0.98, *p* = 0.70).

Revealing behavior-specific differences, univariate ANOVA [factors same as MANOVA] found significant group x time interactions for 4 of the 12 behaviors: freeze (F86,2064 = 2.13, *p* = *1.85 x 10^-8^*; Figure 3C), locomote (F86,2064 = 3.72, *p* = 5.04 x 10^-26^; Figure 3D), scale (F86,2064 = 1.53, *p* = 0.003; Figure 3G), and light rear (F86,2064 = 1.47, *p* = 0.004; Figure 3I). Paired rats showed greater freezing only during the cue period, as well as greater locomotion during the cue and post-cue periods. Unpaired rats showed greater scaling and light rearing during the cue and post-cue periods. Univariate ANOVA found a significant group x time x intensity interaction only for rearing (F86,2064 = 2.32, *p* = 3.87 x 10^-4^). Ethograms separating 0.5 and 0.35 mA rats are shown in Figure 4 – Supplemental Figure 1. Paired rats receiving the 0.5 mA foot shock suppressed rearing during cue illumination while 0.35 mA rats did not.

### Ethograms predict group membership (paired vs. unpaired)

Given the multitude of differing behaviors, it should be possible to use ethogram data to predict group membership. To do this we turned to latent discriminant analysis. We used ethogram data to create a linear array for each of the 64 sessions (32 conditioning and 32 extinction). Linear discriminant analysis sampled a subset of the 64 sessions to train a linear classifier to predict group membership. The linear classifier was then tested on the held-out sessions. This process was repeated 100 times. The data of primary interest was mean ± SEM group classification accuracy for the held-out data, with chance classification being 50%.

We first performed linear discriminant analysis for the total ethogram data (1044 value array for each session, 12 behaviors x 87 samples). We compared linear discriminant analysis of the total ethogram data to two kinds of shuffled ethogram data. Session-shuffled ethogram data were shuffled by row, meaning that the relationship between group membership and ethogram was lost. Temporal-shuffled ethogram data were shuffled by column, meaning the group membership information was intact but the temporal structure of behavior was lost.

Ethograms predicted group membership, and prediction did not deteriorate when temporal structure was lost. Mean ± SEM classification accuracy for the total ethogram data was 82.64 ± 0.003%, exceeding chance (one sample t-test, *p* = 9.31 x 10^-109^; Figure 5A). Mean ± SEM classification accuracy for session-shuffled data was 49.53 ± 0.008%, which did not differ from chance (one sample t-test, *p* = 0.57). Mean ± SEM classification accuracy for the temporal-shuffled ethogram data was 83.16 ± 0.003%, exceeding chance (one sample t-test, *p* = 1.71 x 10^-112^). Total ethogram and temporal-shuffled ethogram classification did not differ from one another (one sample t-test, *p* = 0.18).

**Figure 5.**
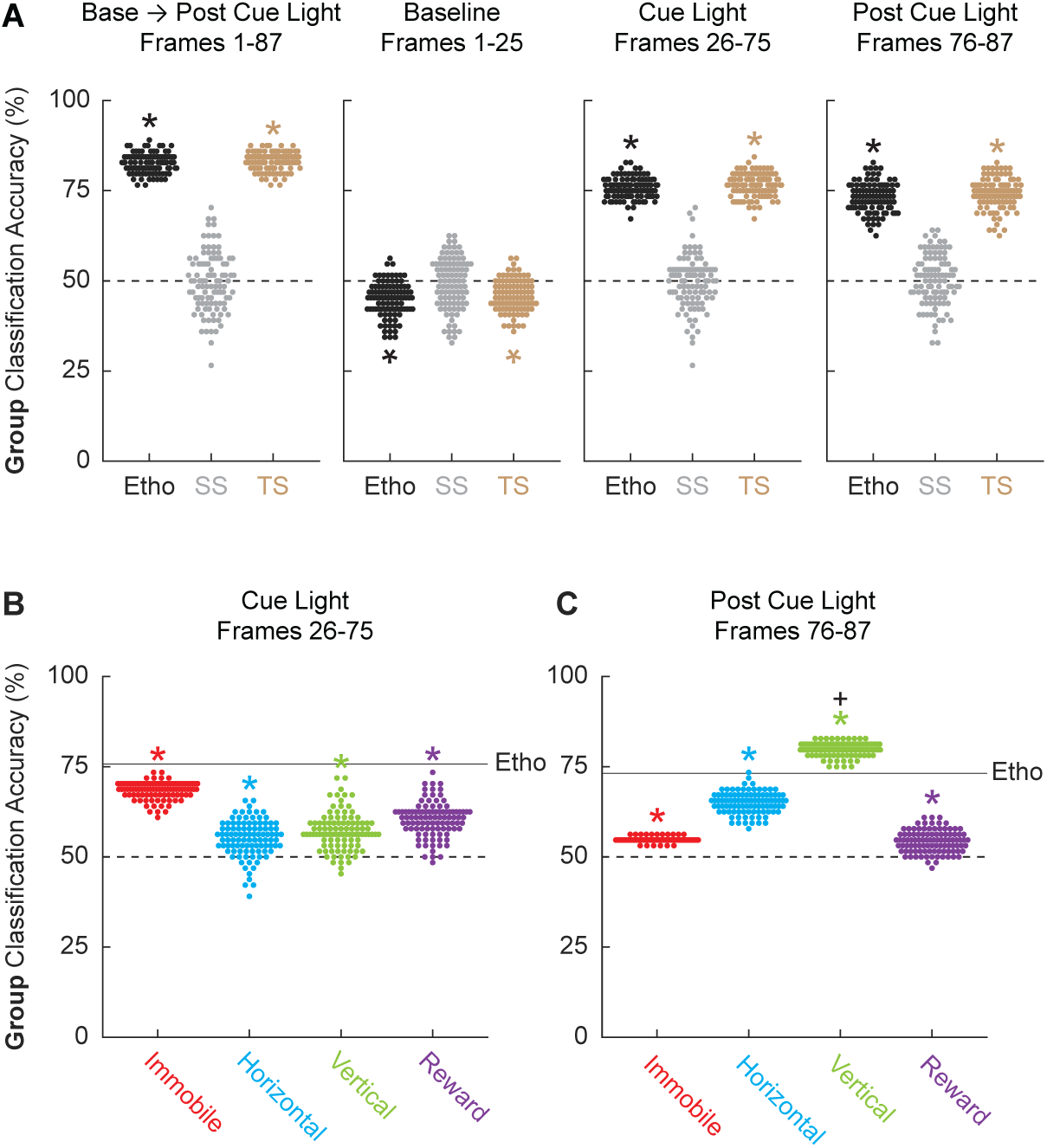
Linear discriminant analysis results. **(A)** Group classification accuracy (paired vs. unpaired) is shown for the total ethogram data (left), baseline (second from left), cue light (second from right), and post cue light (right). Ethogram data were intact (Etho, black), shuffled by session (SS, gray), or shuffled temporally (TS, tan). Each data point represents the accuracy of a single model. The dotted line indicates chance classification. (**B**) Group classification accuracy for the cue light illumination period is shown for separate, multi-behavior categories (Immobile, red; Horizontal, cyan; Vertical, green; and Reward, purple). The dotted line indicates chance classification while the solid line indicates mean classification accuracy for the total ethogram data during cue light illumination. (**C**) Group classification accuracy for the post cue period is shown for separate, multi-behavior categories (color as in B). The dotted line indicates chance classification while the solid line indicates mean classification accuracy for the total ethogram data post cue light illumination. *Significance of a one-sample t-test compared to 0.5 (chance). ^+^Significance of an independent-samples t-test comparing Vertical classification to total ethogram classification.

To determine which trial components provided the most group information we broke the 87 frames into baseline (frames 1-25), cue light illumination (frames 26-75), and post cue light (frames 76-87) periods. Group membership could not be classified from baseline ethogram data. Indeed, group classification was below chance for the total ethogram data (44.91 ± 0.005%; one sample t-test, *p* = 1.94 x 10^-18^) and the temporal-shuffled ethogram data (45.91 ± 0.004%; one sample t-test, *p* = 2.88 x 10^-15^). Group membership was readily classified during the cue light illumination period [total ethogram data (75.75 ± 0.003%; one sample t-test, *p* = 7.91 x 10^-96^); and the temporal-shuffled ethogram data (76.61 ± 0.003%; one sample t-test, *p* = 5.06 x 10^-93^)] as well as the post cue light period [total ethogram data (73.16 ± 0.004%; one sample t-test, *p* = 1.50 x 10^-76^); and the temporal-shuffled ethogram data (73.93 ± 0.004%; one sample t-test, *p* = 2.28 x 10^-77^)]. Observing comparable classification during the cue and post cue periods means that group classification could not be simply attributed to shock responding. In fact, latent discrimination analysis to classify session type (conditioning vs. extinction) from all 87 frames for the total ethogram data achieved chance accuracy (49.55%).

Finally, we asked if comparable group classification could be achieved using only subsets of behavior categories. To do this we divided the ethogram into 4 multi-behavior categories: Immobile (Freeze and Stretch), Horizontal (Locomote and Backpedal), Vertical (Rear, Scale, Jump, Light Rear, and Light Scale), and Reward (Cup and Port). Group classification during cue light illumination was greater chance for each multi-behavior category, but was variable across categories and worse than classification based on the total ethogram: Immobile (68.50 ± 0.002%; one sample t-test compared to 50%, *p* = 5.98 x 10^-91^), Horizontal (55.16 ± 0.005%; one sample t-test compared to 50%, *p* = 2.35 x 10^-17^), Vertical (57.19 ± 0.005%; one sample t-test compared to 50%, *p* = 2.66 x 10^-25^), and Reward (60.44 ± 0.005%; one sample t-test compared to 50%, *p* = 8.40 x 10^-40^). Group classification post cue light illumination was also greater chance for each multi-behavior category, but again was variable across categories and only Vertical classification exceeded the total ethogram: Immobile (54.77 ± 0.0006%; one sample t-test compared to 50%, *p* = 1.47 x 10^-88^), Horizontal (65.06 ± 0.003%; one sample t-test compared to 50%, *p* = 6.35 x 10^-73^), Vertical (79.84 ± 0.002%; one sample t-test compared to 50%, *p* = 9.64 x 10^-121^), and Reward (54.30 ± 0.003%; one sample t-test compared to 50%, *p* = 2.48 x 10^-25^).

## Discussion

We set out to comprehensively quantify behaviors engaged by a shock-paired visual cue. First, we found that the ability of a visual cue to support conditioned suppression depends on foot shock intensity (Holland, 1979). In rats showing robust conditioned suppression of reward-related responding, we also observed robust suppression of scaling and rearing, as well as suppression of light-directed scaling and rearing. Freezing was specific to a shock-paired visual cue, but was shown at low levels during conditioning and even lower levels during extinction testing. Our observation of minimal conditioned freezing, but robust conditioned by a shock-paired visual cue is consistent with prior studies (Kim et al., 1996). Finally, a shock-paired visual cue elicited locomotion during both cues and post-cue periods in conditioning as well as extinction. The results bolster our prior findings of diverse fear conditioned behaviors, and further demonstrate the ability of a shock-paired cue to elicit locomotion (Chu et al., 2024).

Compared to studies using auditory cues (Walker et al., 2018), visual cues produces higher levels of unconditioned suppression of reward-related behavior that persist for more sessions. This is likely because a visual stimulus, beyond illuminating the physical space occupied by the stimulus, provides more general context illumination. Visual cue illumination then provides the basis for two different types of behavior: stimulus-directed orienting and context exploration. This means that a shock-paired visual cue will not only suppress reward-related behavior, but stimulus orienting and context exploration. While it is possible that all three are achieved through response competition with freezing and locomotion (which would also compete with one another), it is more likely the capacity to suppress each of the three is independent of other overt behaviors. Conditioned suppression of contextual exploration would represent a novel means of studying behavioral processes related to agoraphobia.

Our experimental design included the balanced use of female and male rats. This was done primarily for experimental rigor, but also because darting (similar to our locomotion) has been shown to be more prevalent in female rats. In general, we found female and male rats to be much behaviorally similar than different. However, we did find that during conditioning females tended to show longer jumping bouts and less frequent backpedaling bouts following shock presentation. Our finding of differences in shock-responsiveness that do not manifest as differences in cue-elicited behavior are consistent with recent results (Mitchell et al., 2024).

Our high-resolution approach to scoring (5 frames per second) gave us an unprecedented look at the temporal organization of behavior. Looking at ethograms, behavioral transitions across cue presentation, then from cue offset to shock onset were evident. Surprising to us, this temporal organization was not essential to classify rats as paired or unpaired based on their ethogram data. This means that *whether* a behavior occurred during a trial was more telling of paired or unpaired identity than *when* that behavior occurred. Consistent with this description, paired and unpaired identity could be classified above chance from only the freezing and stretching behavior data. This is despite the fact that freezing and stretching accounted for only a very small portion of the data. Still, group classification was most effective when all behaviors were considered across trial entirety.

The temporal organization of locomotion was highly conserved from conditioning to extinction. As expected, paired rats showed robust locomotion following shock delivery during the conditioning sessions, an unconditioned response. Locomotion also occurred during cue light illumination in those same sessions, a conditioned response. Visual-cue elicited locomotion was further evident during the extinction session, again demonstrating a conditioned response. However, not only did we observe locomotion following cue offset during extinction (when no shock is given), but its timing and magnitude strongly resembled that from the response to the shock itself. In fact, the organization of paired behavior during the extinction session was so similar to the conditioning session that linear discriminant analysis performed only on paired rats could not accurately classify conditioning vs. extinction sessions. Not only does a shock-paired visual cue engage diverse behaviors but this capacity transfers very well to an extinction setting.

Studies of Pavlovian fear conditioning measuring freezing have been invaluable for mapping neural circuits for fear learning and expression. These results join a host of recent studies (Le et al., 2023; Chanthongdee et al., 2024; Mitchell et al., 2024) demonstrating an impressive diversity of fear conditioned behaviors. Mapping neural circuits for these diverse fear behaviors will mean a more complete understanding of the brain basis of fear, that is more likely to translate to the full spectrum of anxiety and stressor-related disorders.

## Materials and Methods

### Subjects

Experiments 1 and 2 each used thirty-two adult Long Evans rats (sixteen females), for a total of sixty-four rats (thirty-two females). Rats were obtained from Charles River Laboratories on postnatal day 55. Rats were single-housed on a 12-hr light cycle (lights off at 6:00pm) and maintained at their initial body weight with standard laboratory chow (18% Protein Rodent Diet #2018, Harlan Teklad Global Diets, Madison, WI). Water was available *ad libitum* in the home cage. All experiments were carried out in accordance with the NIH guidelines regarding the care and use of rats for experimental procedures. All procedures were approved by the Boston College Animal Care and Use Committee. The Boston College experimental protocol supporting these procedures is 2024-001.

### Behavior apparatus

The behavior apparatus consisted of eight individual chambers with aluminum front and back walls, clear acrylic sides and top, and a grid floor. LED strips emitting 940 nm light were affixed to the acrylic top to constantly illuminate the behavioral chamber for frame capture. 940 nm illumination was chosen because rats do not detect light wavelengths exceeding 930 nm (Nikbakht and Diamond, 2021). An array of three infrared lights were mounted to the chamber door to track trial period (baseline, light, and post-light). Each grid floor bar was electrically connected to an aversive shock generator (Med Associates, St. Albans, VT). An external food cup, a central port equipped with infrared photocells, and a key light were present on one wall.

### Pellet exposure and nose poke shaping

Identical pellet exposure and nose poke shaping were performed for Experiments 1 and 2. Rats were food-restricted and specifically fed to maintain their body weight throughout behavioral testing. Each rat was given four grams of experimental pellets in their home cage in order to overcome neophobia. Next, the central port was removed from the experimental chamber, and rats received a 30-minute session in which one pellet was delivered every minute. The central port was returned to the experimental chamber for the remainder of behavioral testing. Each rat was then shaped to nose poke in the central port for experimental pellet delivery using a fixed ratio schedule in which one nose poke into the port yielded one pellet. Shaping sessions lasted 30 min or until approximately 50 nose pokes were completed. Each rat then received five sessions during which nose pokes into the port were reinforced on a variable interval schedule. Session 1 used a variable interval 30 s schedule (poking into the port was reinforced every 30 s on average). All remaining sessions used a variable interval 60 s schedule. For the remainder of behavioral testing, nose pokes were reinforced on a variable interval 60 s schedule independent of light and shock presentation.

### Light pre-exposure

Identical light pre-exposure was performed for Experiments 1 and 2. Each rat was pre-exposed to the 10-s light, four times during two, 43-minute sessions (Monday and Tuesday). The two pre-exposure sessions meant that each rat received eight total, light pre-exposures. Although light intensity was greatest at the source, light presentation generally illuminated the experimental chamber which was completely dark to the rat at all other times.

### Light and shock presentations

Following pre-exposure, the 32 rats in each experiment were evenly divided into two groups: Paired (16 rats, 8 females) and Unpaired (16 rats, 8 females). Each Paired and Unpaired rat received 4 light, and 4 shock presentations during two, 43-minute sessions (Wednesday and Thursday). The two sessions meant that each rat received 8 total light and shock presentations. For rats in the Paired group, the light was presented for 10 s. Light offset precisely coincided with foot shock presentation (0.5 s). For rats in the Unpaired group, light (10 s) and shock (0.5 s) were presented at distant times. The inter-event interval (in which both light and shock are considered events) was greater than five minutes. The order of Unpaired light and shock presentation was randomly determined and differed for each rat, each session.

### Foot shock intensity

For both experiments, two levels of foot shock intensity were used. For Experiment 1, half of the rats received a 0.25 mA foot shock, and half 0.15 mA foot shock. The total experimental design yield 4 groups: Paired 0.25 (8 rats, 4 females), Paired 0.15 (8 rats, 4 females), Unpaired 0.25 (8 rats, 4 females), and Unpaired 0.15 (8 rats, 4 females). For Experiment 2, half of the rats received a 0.50 mA foot shock, and half 0.35 mA foot shock. The total experimental design yield 4 groups: Paired 0.50 (8 rats, 4 females), Paired 0.35 (8 rats, 4 females), Unpaired 0.50 (8 rats, 4 females), and Unpaired 0.35 (8 rats, 4 females).

### Extinction test

Identical extinction tests were performed for Experiments 1 and 2. The single, extinction session (Friday) was identical to the pre-exposure session. Each rat received four, 10-s light presentations in a 43-minute session.

### Calculating suppression ratio

Time stamps for cue light illumination, shock presentation, and nose pokes (photobeam break) were automatically recorded by the Med Associates program. Baseline nose poke rate was calculated for each trial by counting the number of pokes during the 20-s pre-light period and multiplying by 3. Light nose poke rate was calculated for each trial by counting the number of pokes during the 10-s light period and multiplying by 6. Nose poke suppression was calculated as a ratio: (baseline poke rate – light poke rate) / (baseline poke rate + light poke rate). A suppression ratio of ‘1’ indicated complete suppression of nose poking during light presentation relative to baseline. A suppression ratio of indicated ‘0’ indicates equivalent nose poke rates during baseline and light presentation. Gradations in suppression ratio between 1 and 0 indicated intermediate levels of nose poke suppression during light presentation relative to baseline. Negative suppression ratios indicated increased nose poke rates during light presentation relative to baseline.

### Frame capture system

Behavior frames were captured using Imaging Source monochrome cameras (DMK 37BUX28; USB 3.1, 1/2.9“ Sony Pregius IMX287, global shutter, resolution 720x540, trigger in, digital out, C/CS-mount). Frame capture was triggered by the Med Associates behavior program. The 28V Med Associates pulse was converted to a 5V TTL pulse via Adapter (SG-231, Med Associates, St. Albans, VT). The TTL adapter was wired to the camera’s trigger input. Captured frames were saved to a PC (OptiPlex 7470 All-in-One) running IC Capture software (Imaging Source). Frame capture began precisely 5 s before light onset, continued throughout 10-s light presentation, and ended 2.5 s following light offset. For Paired rats this meant frames were captured during the 0.5 s shock and the 2 s following. For Unpaired rats, the empty 2.5 s period was captured. Frames were captured at a rate of 5 per second, with a target of capturing 87 frames per trial, and 348 frames per session.

### Anonymizing trial information

A total of 256 trials were scored from the 32 rats in Experiment 2. Trials came from the second session of light/shock presentations (Thursday) and the extinction test (Friday). We anonymized trial information in order to score behavior without bias. The numerical information from each trial (session #, rat # and trial #) was encrypted as a unique number sequence. A unique word was prepended to the sequence. The result was that each trial was converted into a unique word+number sequence. For example, trial dw01_01_03 (rat #1, session #1 and trial #3) would be encrypted as: abundant28515581. The trials were randomly assigned to 6 observers. The result of trial anonymization was that observers were completely blind to subject, group, and session. Further, random assignment meant that the 4 trials composing a single session were scored by different observers.

### Post-acquisition frame processing

A Matlab script sorted the 348 frames into 4 folders, one for each trial, each containing 87 frames. Each 87-frame trial was made into an 87-slide PowerPoint presentation to be used for hand scoring.

### Behavior categories and definitions

Frames were scored as one of twelve mutually exclusive behavior categories, defined as follows:

<u>Background</u>. Specific behavior cannot be discerned because the rat is turned away from the camera or position of forepaws is not clear, or because the rat is not engaged in any of the other behaviors.

<u>Backpedal</u>. Rapid backward displacement of the body. Often (not always) the head is down and/or back is arched. The backward component of movement must be stronger than the lateral component. A lateral movement is not a backpedal. Backpedal considers the current frame (t) and the next two frames (t+1 and t+2). By frame t+2 the body and both back feet must be displaced backward relative to frame t. The rat can move the body and both feet by t+1, t+2 or move in combination of both frames; all count as backpedal for trial t. (See illustration for reference)

<u>Cup</u>. Any part of the nose above the food cup but below the nose port.

<u>Freeze</u>. Arched back and stiff, rigid posture in the absence of movement, all four limbs on the floor (often accompanied by hyperventilation and piloerection). Side to side head movements and up and down head movements that do not disturb rigid posture are permitted. Activity such as sniffing, or investigation of the bars is not freezing. Freezing, as opposed to pausing, is likely to be 3 or more frames (600+ ms) long.

<u>Groom</u>. Any scratching, licking, or washing of the body.

<u>Jump</u>. All four limbs off the floor. Includes hanging which is distinguished when hind legs are hanging freely.

<u>Locomote</u>. Propelling body across chamber on all four feet, as defined by movement of back feet. Movement of back feet with front feet off floor is rearing. Locomote considers the current frame (t) and the next two frames (t+1 and t+2). By frame t+2 the body and both back feet must be displaced forward relative to frame t. The rat can move the body and both feet by t+1, t+2 or move in combination of both frames; all count as locomote for trial t.

<u>Port</u>. Any part of the nose in the port. Often standing still in front of the port but sometimes tilting head sideways with the body off to the side of the port.

<u>Rear</u>. One or two hind legs on the grid floor with both forepaws *o*ff the grid floor and not on the food cup. Usually (not always) stretching to full extent, forepaws usually (not always) on top of side walls of the chamber, often pawing walls; may be accompanied by sniffing or slow side-to-side movement of head. Does not include grooming movements or eating, even if performed while standing on hind legs.

<u>Light rear</u>. A rear happening within the 3D space defined by these planes: nose is above the poke (X), between the metal columns in the light area (Y), within the depth of the front edge of the food cup (Z). If the rats body is outside of the columns BUT the nose is within the defined 3D space, it is a Light Rear. Often the body will be near the wall and/or the forepaws will be on the wall. The nose can be above the light. The light does not have to be on to be scored as a Light Rear.

<u>Scale</u>. All four limbs off the floor but at least two limbs on the side of the chamber. Standing on the food cup counts as scaling.

<u>Light scale</u>. A scale that is happening when the nose is above the poke and between the metal columns in the light area. Including above the light. The light does not have to be on to be scored as a light scale.

<u>Stretch</u>. The body is elongated with the back posture ’flatter’ than normal. Stretching is often accompanied by immobility, like freezing, but is distinguished by the shape of the back.

### Frame scoring system

Frames were scored using a specific procedure. Frames were first watched in real time in Microsoft PowerPoint by setting the slide duration and transition to 0.19 s, then playing as a slideshow. Behaviors clearly observed were noted. Next, the observer went through all the frames scoring one behavior at a time. A standard scoring sequence was used: groom, port, cup, light rear, rear, light scale, scale, jump, freeze, locomote, backpedal, stretch. When the specific behavior was observed in a frame, that frame was labeled. Once all behaviors had been scored, the video was re-watched for freezing. The unlabeled frames were then labeled ‘background’. Finally, all background frames were checked to ensure they did not contain a defined behavior.

### Statistical analyses

Multivariate analysis of variance (MANOVA) and univariate analysis of variance (ANOVA) were performed for suppression ratios, and specific behaviors. Sex was used as a factor for all analyses. Group, session, intensity, and time were used as factors when relevant. Univariate ANOVA following MANOVA used a Bonferroni-corrected *p* value significance of 0.004167 (0.05/12) to account for the twelve quantified behaviors. Post-hoc comparisons were made using independent samples and one-sample t-tests.

### Linear discriminant analysis

Linear discriminant analysis (MATLAB’s fitcdiscr function) was used to predict group membership (paired vs. unpaired) from ethogram data. LDA is a linear classification method that projects the ethogram features into a space that maximizes the separation between groups.

Each LDA model used the entire dataset without any initial data splitting. To evaluate the performance of the LDA model, 10-fold cross-validation was applied using MATLAB’s crossval function. Cross-validation divides the dataset into 10 equal parts (folds). For each iteration, the model was trained on 9 folds and tested on the remaining fold. This process was repeated 10 times so that each fold served as a test set once. The accuracy of the model (defined as 1 minus the loss, 1-L) was assessed by computing the classification error for each fold, and the overall performance was averaged across all folds. The crossval function was used in conjunction with kfoldLoss, which returns the average classification error across the folds. The performance of the LDA model was quantified using the average classification accuracy across the 10 folds. This approach provided a robust estimate of the model’s generalization capability to unseen data, reducing the likelihood of overfitting. One hundred LDA models were made for each analyses. Prediction accuracies from each model were saved and used to calculate mean ± SEM prediction accuracies across all iterations, for each analysis.

#### Box 1. Behavior definitions

<u>Background</u>. Specific behavior cannot be discerned because the rat is turned away from the camera or position of forepaws is not clear, or because the rat is not engaged in any of the other behaviors.

<u>Backpedal</u>. Rapid backward displacement of the body. Often (not always) the head is down and/or back is arched. The backward component of movement must be stronger than the lateral component. A lateral movement is not a backpedal. Backpedal considers the current frame (t) and the next two frames (t+1 and t+2). By frame t+2 the body and both back feet must be displaced backward relative to frame t. The rat can move the body and both feet by t+1, t+2 or move in combination of both frames; all count as backpedal for trial t. (See illustration for reference)

<u>Cup</u>. Any part of the nose above the food cup but below the nose port.

<u>Freeze</u>. Arched back and stiff, rigid posture in the absence of movement, all four limbs on the floor (often accompanied by hyperventilation and piloerection). Side to side head movements and up and down head movements that do not disturb rigid posture are permitted. Activity such as sniffing, or investigation of the bars is not freezing. Freezing, as opposed to pausing, is likely to be 3 or more frames (600+ ms) long.

<u>Groom</u>. Any scratching, licking, or washing of the body.

<u>Jump</u>. All four limbs off the floor. Includes hanging which is distinguished when hind legs are hanging freely.

<u>Locomote</u>. Propelling body across chamber on all four feet, as defined by movement of back feet. Movement of back feet with front feet off floor is rearing. Locomote considers the current frame (t) and the next two frames (t+1 and t+2). By frame t+2 the body and both back feet must be displaced forward relative to frame t. The rat can move the body and both feet by t+1, t+2 or move in combination of both frames; all count as locomote for trial t.

<u>Port</u>. Any part of the nose in the port. Often standing still in front of the port but sometimes tilting head sideways with the body off to the side of the port.

<u>Rear</u>. One or two hind legs on the grid floor with both forepaws *o*ff the grid floor and not on the food cup. Usually (not always) stretching to full extent, forepaws usually (not always) on top of side walls of the chamber, often pawing walls; may be accompanied by sniffing or slow side-to-side movement of head. Does not include grooming movements or eating, even if performed while standing on hind legs.

<u>Light rear</u>. A rear happening within the 3D space defined by these planes: nose is above the poke (X), between the metal columns in the light area (Y), within the depth of the front edge of the food cup (Z). If the rats body is outside of the columns BUT the nose is within the defined 3D space, it is a Light Rear. Often the body will be near the wall and/or the forepaws will be on the wall. The nose can be above the light. The light does not have to be on to be scored as a Light Rear.

<u>Scale</u>. All four limbs off the floor but at least two limbs on the side of the chamber. Standing on the food cup counts as scaling.

<u>Light scale</u>. A scale that is happening when the nose is above the poke and between the metal columns in the light area. Including above the light. The light does not have to be on to be scored as a light scale.

<u>Stretch</u>. The body is elongated with the back posture ’flatter’ than normal. Stretching is often accompanied by immobility, like freezing, but is distinguished by the shape of the back.

---

**Figure 3 - Supplemental Figure 1.**
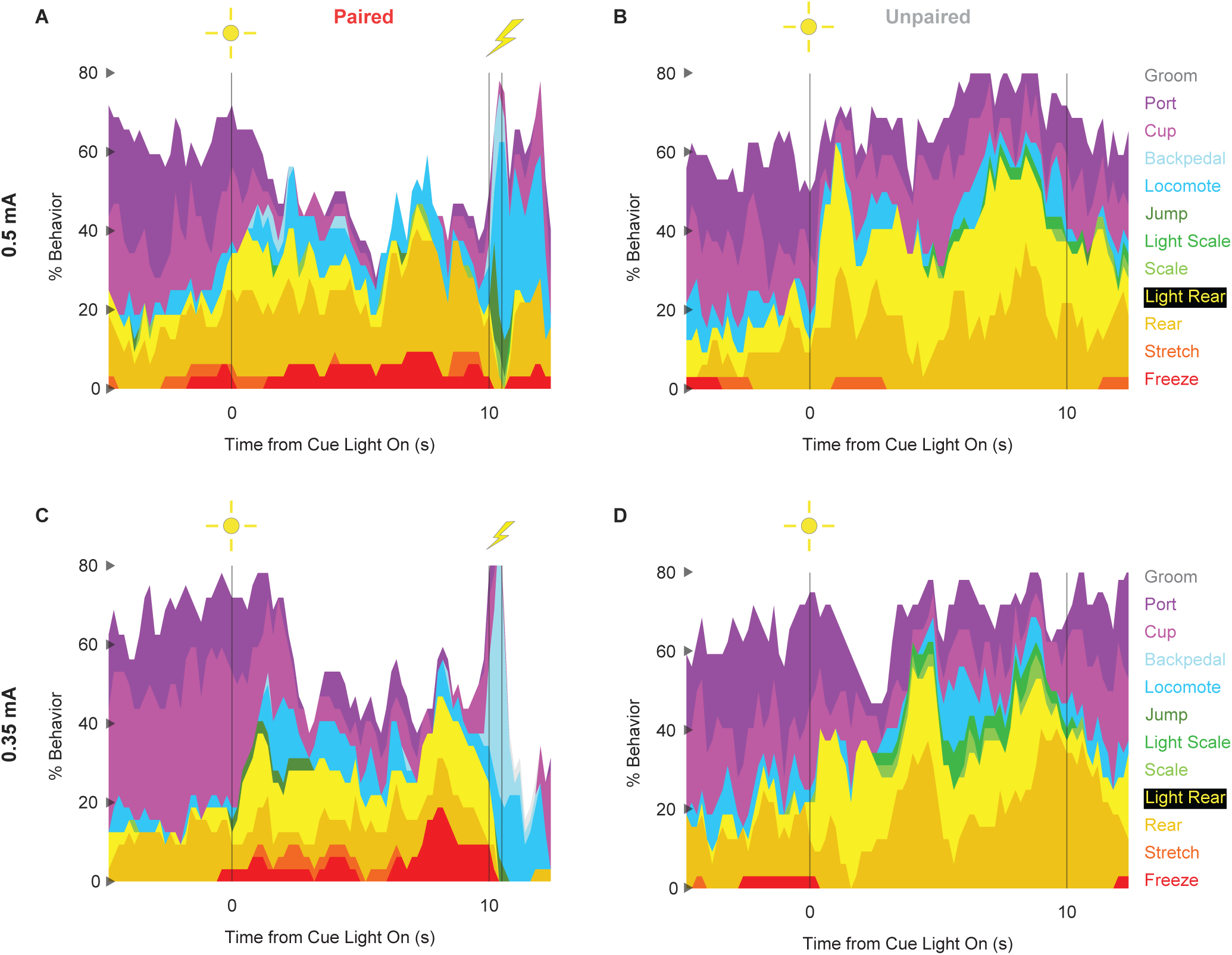
Conditioning ethograms by intensity. (**A**) Ethogram for paired rats receiving the 0.5 mA foot shock shows % behavior (y axis) for each behavioral category in 200-ms intervals from 5-s prior to cue onset (time 0) to 2.5 s following cue offset (time 10). Paired rats had shock delivered at cue offset. Colors for each behavior category: Freeze (red), Stretch (orange), Rear (mustard), Light Rear (yellow), Scale (light green), Light Scale (green), Jump (dark green), Locomote (cyan), Backpedal (sky blue), Cup (magenta), Port (purple), and Groom (gray). (**B**) Ethogram for unpaired rats receiving the 0.5 mA foot shock, details identical to paired rats. Unpaired rats did not receive shock at cue offset. (**C** & **D**) Ethogram formatting is identical to A & B but for paired and unpaired rats receiving the 0.35 mA foot shock.

**Figure 3 - Supplemental Figure 2.**
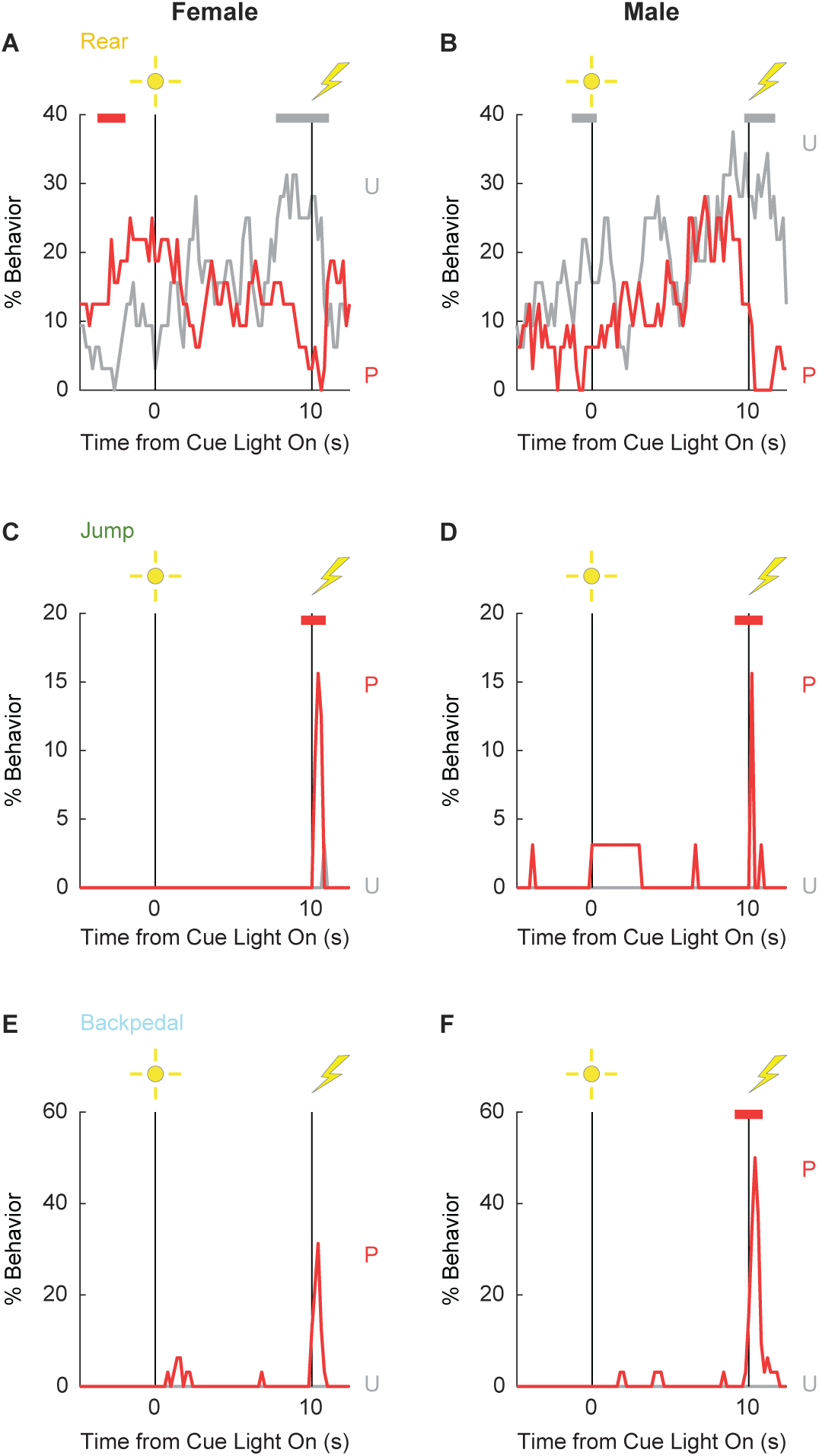
Conditioning line graphs by sex. Line graphs for the 3 behaviors showing a significant group x time x sex interaction are shown: Rear (**A** & **B**), Jump (**C** & **D**), and Backpedal (**E** & **F**). Females are plotted in the left column and males in the right column. % Behavior is plotted in 200-ms intervals from 5-s prior to cue onset (time 0) to 2.5 s following cue offset (time 10). Paired rats (P, red) and unpaired rats (U, gray). Colored bars at the top of each axis indicate 1-s time periods in which paired and unpaired % behavior differed (indepen-dent samples t-test, p < 0.05). Red bars indicate greater behavior in paired rats while gray bars indicate greater behavior in unpaired rats.

**Figure 4 - Supplemental Figure 1.**
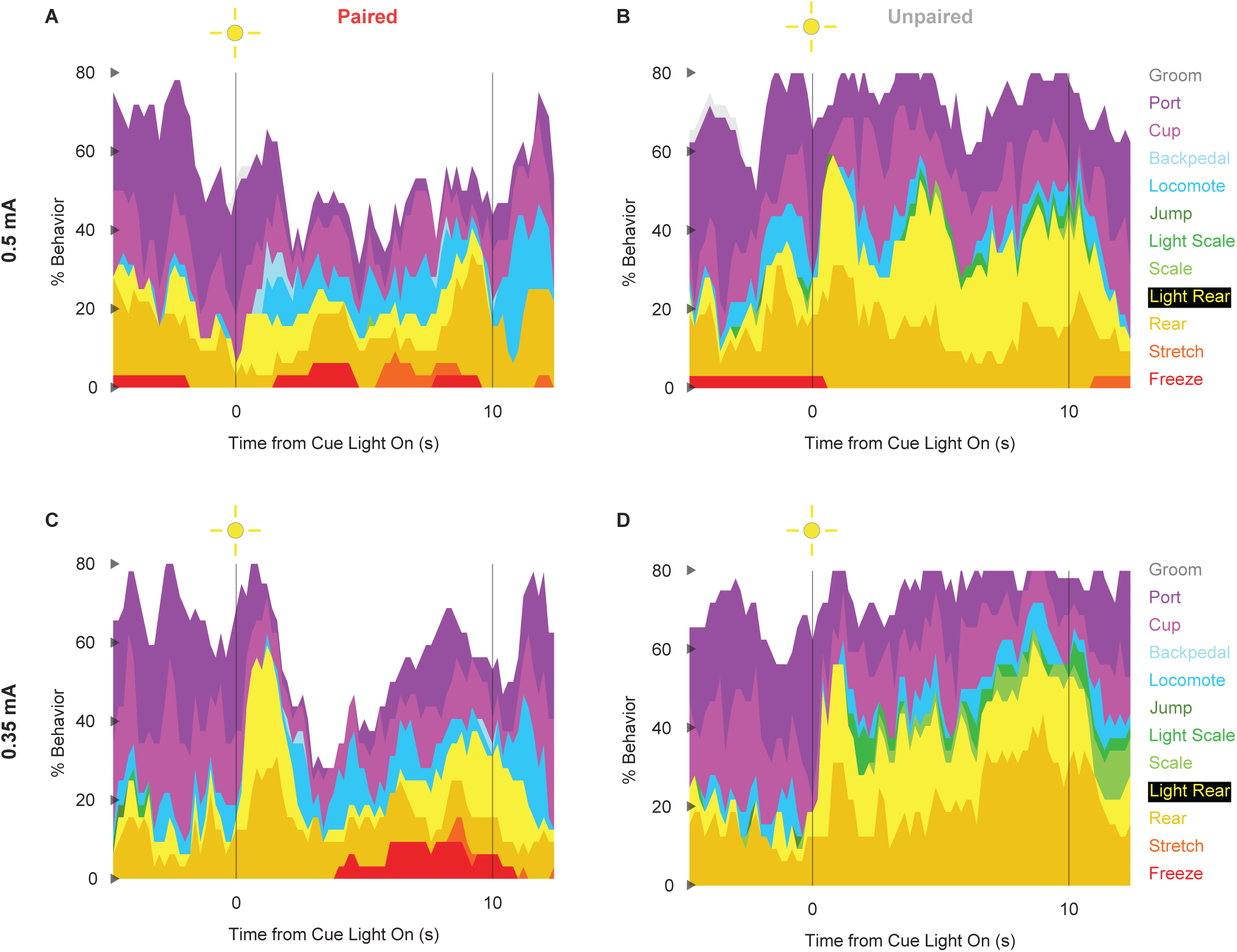
Extinction ethograms by intensity. (**A**) Ethogram for paired rats receiving the 0.5 mA foot shock shows % behavior (y axis) for each behavioral category in 200-ms intervals from 5-s prior to cue onset (time 0) to 2.5 s following cue offset (time 10). Paired rats had shock delivered at cue offset. Colors for each behavior category: Freeze (red), Stretch (orange), Rear (mustard), Light Rear (yellow), Scale (light green), Light Scale (green), Jump (dark green), Locomote (cyan), Backpedal (sky blue), Cup (magenta), Port (purple), and Groom (gray). (**B**) Ethogram for unpaired rats receiving the 0.5 mA foot shock, details identical to paired rats. Unpaired rats did not receive shock at cue offset. (**C** & **D**) Ethogram formatting is identical to A & B but for paired and unpaired rats receiving the 0.35 mA foot shock.

## Literature Cited

Amorapanth P, Nader K, LeDoux JE (1999) Lesions of periaqueductal gray dissociate-conditioned freezing from conditioned suppression behavior in rats. Learning & Memory 6:491–499.

Bolles RC, Collier AC (1976) The effect of predictive cues on freezing in rats. Animal Learning & Behavior 4:6–8.

Borkar CD, Stelly CE, Fu X, Dorofeikova M, Le Q-SE, Vutukuri R, Vo C, Walker A, Basavanhalli S, Duong A, Bean E, Resendez A, Parker JG, Tasker JG, Fadok JP (2024) Top-down control of flight by a non-canonical cortico-amygdala pathway. Nature 625:743–749.

Bouton ME, Bolles RC (1980) Conditioned fear assessed by freezing and by the suppression of three different baselines. Animal Learning & Behavior 8:429–434.

Chanthongdee K, Fuentealba Y, Wahlestedt T, Foulhac L, Kardash T, Coppola A, Heilig M, Barbier E (2024) Comprehensive ethological analysis of fear expression in rats using DeepLabCut and SimBA machine learning model. Front Behav Neurosci 18 Available at: https://www.frontiersin.org/journals/behavioral-neuroscience/articles/10.3389/fnbeh.2024.1440601/full [Accessed August 25, 2024].

Chu A, Gordon NT, DuBois AM, Michel CB, Hanrahan KE, Williams DC, Anzellotti S, McDannald MA (2024) A fear conditioned cue orchestrates a suite of behaviors in rats Hill MN, Wassum KM, Maren S, eds. eLife 13:e82497.

De Oca BM, DeCola JP, Maren S, Fanselow MS (1998) Distinct regions of the periaqueductal gray are involved in the acquisition and expression of defensive responses. Journal of Neuroscience 18:3426–3432.

Estes KW, Skinner BF (1941) Some Quantitative Properties of Anxiety. Journal of Experimental Psychology 29:390–400.

Goosens KA, Maren S (2001) Contextual and auditory fear conditioning are mediated by the lateral, basal, and central amygdaloid nuclei in rats. Learning & Memory 8:148–155.

Gruene TM, Flick K, Stefano A, Shea SD, Shansky RM (2015) Sexually divergent expression of active and passive conditioned fear responses in rats. Elife 4 Available at: ://WOS:000373851900001.

Holland PC (1979) The effects of qualitative and quantitative variation in the US on individual components of Pavlovian appetitive conditioned behavior in rats. Animal Learning & Behavior 7:424–432.

Kim SD, Rivers S, Bevins RA, Ayres JJ (1996) Conditioned stimulus determinants of conditioned response form in Pavlovian fear conditioning. Journal of Experimental Psychology: Animal Behavior Processes 22:87–104.

Le Q-SE, Hereford D, Borkar CD, Aldaco Z, Klar J, Resendez A, Fadok JP (2023) Contributions of associative and non-associative learning to the dynamics of defensive ethograms. eLife 12 Available at: https://elifesciences.org/reviewed-preprints/90414 [Accessed August 21, 2024].

LeDoux JE, Iwata J, Cicchetti P, Reis DJ (1988) Different projections of the central amygdaloid nucleus mediate autonomic and behavioral correlates of conditioned fear. Journal of Neuroscience 8:2517–2529.

Lee JLC, Dickinson A, Everitt BJ (2005) Conditioned suppression and freezing as measures of aversive Pavlovian conditioning: effects of discrete amygdala lesions and overtraining. Behavioural Brain Research 159:221–233.

Mast M, Blanchard RJ, Blanchard DC (1982) The relationship between freezing and response suppression in a CER situation. Psychological Record 32:151–167.

McDannald MA (2010) Contributions of the amygdala central nucleus and ventrolateral periaqueductal grey to freezing and instrumental suppression in Pavlovian fear conditioning. Behavioural Brain Research 211:111–117.

McDannald MA (2023) Pavlovian Fear Conditioning Is More than You Think It Is. J Neurosci 43:8079–8087.

McDannald MA, Galarce EM (2011) Measuring Pavlovian fear with conditioned freezing and conditioned suppression reveals different roles for the basolateral amygdala. Brain Research 1374:82–89.

Mitchell JR, Trettel SG, Li AJ, Wasielewski S, Huckleberry KA, Fanikos M, Golden E, Laine MA, Shansky RM (2022) Darting across space and time: parametric modulators of sex-biased conditioned fear responses. Learn Mem 29:171–180.

Mitchell JR, Vincelette L, Tuberman S, Sheppard V, Bergeron E, Calitri R, Clark R, Cody C, Kannan A, Keith J, Parakoyi A, Pikus M, Vance V, Ziane L, Brenhouse H, Laine MA, Shansky RM (2024) Behavioral and neural correlates of diverse conditioned fear responses in male and female rats. :2024.08.20.608817 Available at: https://www.biorxiv.org/content/10.1101/2024.08.20.608817v1 [Accessed August 25, 2024].

Totty MS, Warren N, Huddleston I, Ramanathan KR, Ressler RL, Oleksiak CR, Maren S (2021) Behavioral and brain mechanisms mediating conditioned flight behavior in rats. Sci Rep 11:8215.

Trott JM, Hoffman AN, Zhuravka I, Fanselow MS (2022) Conditional and unconditional components of aversively motivated freezing, flight and darting in mice Penzo M, Taffe MA, Penzo M, Cain C, eds. eLife 11:e75663.

Walker RA, Andreansky C, Ray MH, McDannald MA (2018) Early adolescent adversity inflates threat estimation in females and promotes alcohol use initiation in both sexes. Behav Neurosci 132:171–182.

